# Phosphorothioate-modified DNA oligonucleotides inactivate CRISPR-Cpf1 mediated genome editing

**DOI:** 10.1101/253757

**Authors:** Bin Li, Chunxi Zeng, Wenqing Li, Xinfu Zhang, Xiao Luo, Weiyu Zhao, Chengxiang Zhang, Yizhou Dong

## Abstract

CRISPR-Cpf1, a microbial adaptive immune system discovered from *Prevotella* and *Francisella* 1, employs a single-stranded CRISPR RNA (crRNA) to induce double stranded DNA breaks^1^. To modulate genome editing activity of Cpf1 in human cells, we designed a series of crRNA variants including DNA-crRNA and RNA-crRNA duplexes, and identified that phosphorothioate (PS)-modified DNA-crRNA duplex completely blocked the function of Cpf1 mediated gene editing. More importantly, without prehybridization, this PS-modified DNA was able to regulate Cpf1 activity in a time-and dose-dependent manner. Mechanistic studies indicate that PS-modified DNA oligonucleotides hinder the binding between Cpf1-crRNA complex and target DNA substrate. Consequently, phosphorothioate-modified DNA oligonucleotides provide a tunable platform to inactivate Cpf1 mediated genome editing.

---

Cpf1 (CRISPR from *Prevotella* and *Francisella* 1) is one of the bacterial endonucleases that can induce double stranded DNA breaks under the guidance of a single CRISPR RNA (crRNA)^1^. Among the Cpf1 orthologs, AsCpf1 (Cpf1 from *A cidaminococcussp.)* and LbCpf1 (Cpf1 from *Lachnospiraceae bacterium)* were reported for genome editing in human cells^1^. The wild-type crRNA of CRISPR-Cpf1 system comprises a 5’-handle engaging Cpf1 recognition and a guide segment interacting with target DNA sequences through complementary bindings ^1-3^. Crystal structure of the Cpf1-crRNA-dsDNA complex uncovers the T-rich PAM recognition and cleavage mechanism by Cpf1^2,4-7.^ Based on its unique genome editing properties, the CRISPR-Cpf1 system has recently been applied in diverse eukaryotic species including plants and animals to achieve genome editing^8-16^.

Although the CRIPSR system offers a powerful platform for genome editing, a number of challenges exist for its therapeutic applications including genome editing efficiency and potential side effects ^17^. Previously, extensive efforts have been made to improve genome editing efficiency^2,18-21,^. Meanwhile, researchers investigated numerous approaches to modulate the activity of the CRISPR system. Especially, when severe side effects occur in therapeutic applications using this system, it is essential to prepare an effective and fast mechanism to switch off its function. Recently, anti-CRISPR proteins from bacteriophage or bacteria were discovered to inhibit the function of *Neisseria meningitidis* Cas9^22^, *Listeria monocytogenes* Cas9^23^ or *Streptococcus pyogenes* Cas9^23,24.^ In addition, multiple strategies such as chemical-, temperature-and light-controlled approaches were developed to trigger the CRISPR-Cas9 system^25,26.^ Herein, we investigated an array of crRNA duplexes (crDuplex) hybridized with DNA or RNA oligonucleotides, and identified that PS-modified DNA oligonucleotides complementary to crRNA was capable of switching off the activity of CRISPR-Cpf1 system.

## Results

### Effects of crRNA duplexes on Cpf1-mediated genome editing

Oligonucleotides are short single stranded DNA or RNA molecules which have been widely used for diverse applications^27,28.^ Previously, chemically modified nucleotides were extensively applied to tune the properties of oligonucleotides including chemical stability and duplex interactions ^21,27-30^. Several representative examples are listed in Figure 1a. Given the crucial role of crRNA in the formation of Cpf1-crRNA-DNA complex, we hypothesized that an oligonucleotide complementary to crRNA may provide a strategy to regulate Cpf1-mediated genome editing. To test this hypothesis, we designed a series of oligonucleotides using unmodified and chemically modified nucleotides (Fig. 1a). Since the crRNA of Cpf1 is composed of a 5’ handle, seed region, and 3’ end^2^, we synthesized oligonucleotides complementary to different regions (5’handle, seed region, 3’ end, or whole sequence *et al)* of the crRNA (Supplementary Fig. 1). We then generated their corresponding crRNA duplexes (termed crDuplex, Supplementary Fig. 1) by annealing an equimolar of oligonucleotides with the crRNA.

**Fig. 1.**
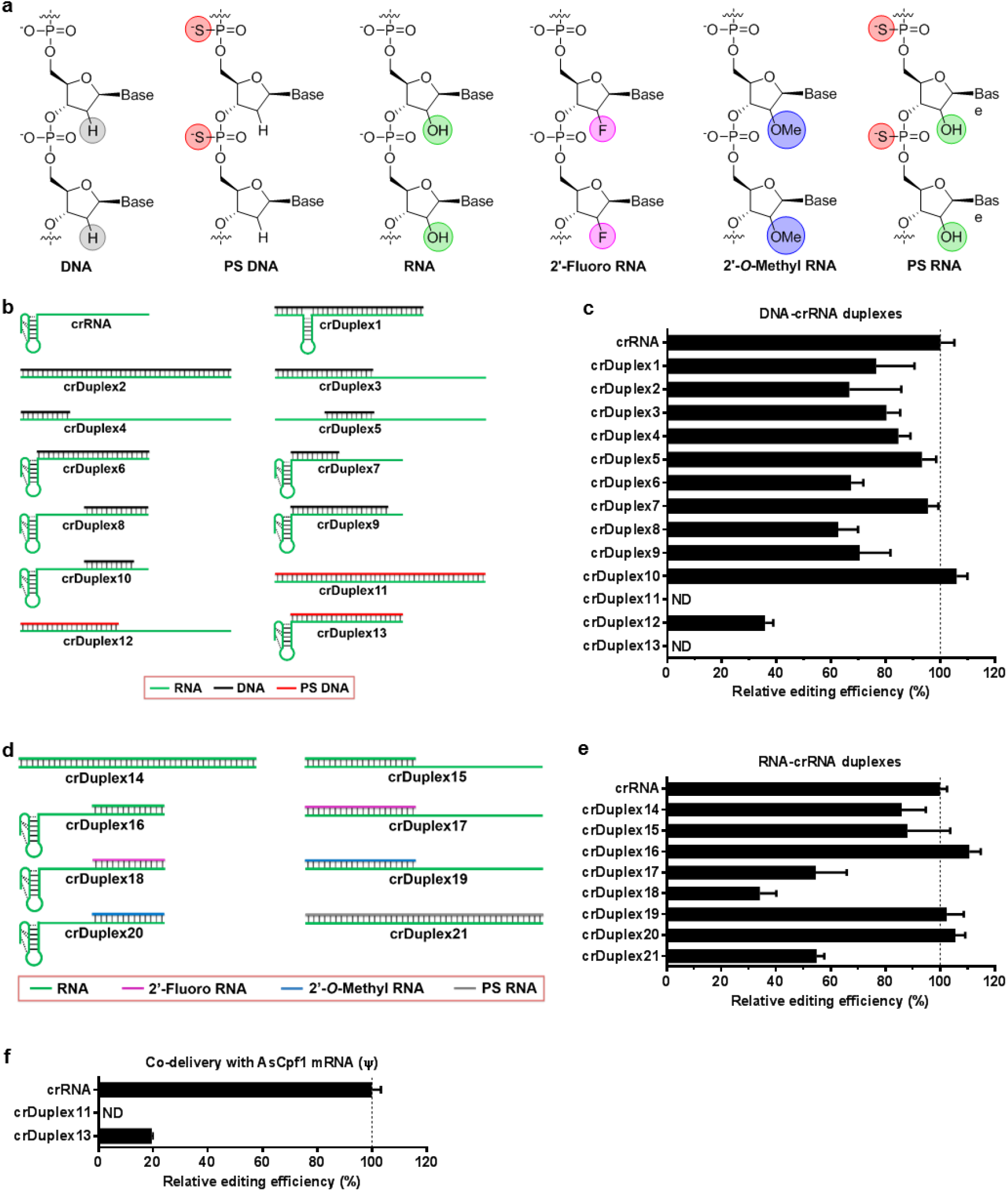
Genome editing efficiency of crRNA duplexes in human cells. **a** Structures of unmodified and chemically modified nucleotides used in this study. **b** Schematic illustration of crRNA and DNA-crRNA duplexes. crRNA consists of a handle (pseudoknot structure) and a guide segment. crRNA duplexes were formed by hybridization of different lengths of unmodified (black) or PS-modified (red) DNA oligonucleotides with various regions of the crRNA. **c** Relative genome editing efficiency of DNA-crRNA duplexes at the *DNMT1* gene locus in the presence of AsCpf1 plasmid in 293T cells. d Schematic illustration of RNA-crRNA duplexes. Duplexes were formed by hybridization of unmodified (green), 2’-fluoro (violet) or 2*’-O*-Methyl modified (blue) RNA oligonucleotides with the crRNA. **e** Relative genome editing efficiency of RNA-crRNA duplexes at the *DNMT1* gene locus in the presence of AsCpf1 plasmid in 293T cells. **f** Relative genome editing efficiency of PS-DNA-crRNA duplexes at the *DNMT1* gene locus in the presence of AsCpf1 mRNA in 293T cells. In all cases, relative genome editing efficiency (%) was determined by the T7E1 cleavage assay, and normalized to that of the wild-type crRNA group. All data are expressed as the mean ± s.d. from three biological replicates. ND, not detectable.

As shown in Figure 1b, the first type of crDuplex, DNA-crRNA, were constructed by hybridization of unmodified or phosphorothioate (PS)-modified DNA oligonucleotide (Fig. 1a) with crRNA at different regions (crDuplex1 to crDuplex13). In the presence of plasmid encoding AsCpf1, crDuplex1 to crDuplex10 (unmodified DNA-crRNA duplexes) retain 60% or more genome editing activity of wild-type crRNA at the *DNMT1* locus in 293T cells. However, we observed no significant correlation between genome editing efficiency and the hybrid region, nor genome editing efficiency and the length of oligonucleotide tested (Fig. 1c). Interestingly, PS-modified DNA-crRNA duplexes including crDuplex11, crDuplex12, and crDuplex13 dramatically reduced genome editing efficiency under the same condition, (Fig. 1c). Moreover, crDuplex11 and crDuplex13 completely blocked the function of Cpf1 (Fig. 1c).

DNA and RNA oligonucleotides possess important chemical difference at the 2’-position on the ribose (Fig. 1a). We then examined the second type of crDuplex, RNA-crRNA, in order to explore the effects of RNA oligonucleotides on Cpf1-mediated genome editing. In this study, unmodified, 2’-fluoro, 2’-0-methyl, and PS-modified RNA oligonucleotides (Fig. 1a) were used to produce unmodified RNA-crRNA (crDuplex14-16), 2’-fluoro RNA-crRNA (crDuplex17 and crDuplex18), 2’-O-methyl RNA-crRNA (crDuplex19 and crDuplex20), and PS-modified RNA-crRNA (crDuplex21) (Fig. 1d). As shown in Figure 1e, crDuplex14-16, crDuplex19 and crDuplex20 showed comparable or slightly higher genome editing efficiency in comparison to crRNA, while crDuplex17, crDuplex18, and crDuplex21 reduced cleavage activity at different extent, depending on the pattern of modification (Fig. 1e). We noticed that 2’-fluoro RNA-crRNA (crDuplex17 and crDuplex18) inhibited genome editing activity more potently than unmodified RNA-crRNA (crDuplex14-16) and 2*’-O*-methyl RNA-crRNA (crDuplex19 and crDuplex20). PS-modified RNA-crRNA (crDuplex21) also displayed inhibition activity, which is comparable to 2’-fluoro RNA-crRNA (crDuplex17 and crDuplex18) but not as potent as its corresponding DNA duplex, crDuplex11.

Taken together, these findings suggested that genome editing activity of the CRISPR-Cpf1 system can be affected by oligonucleotides with different chemical modifications, length of DNA or RNA oligonucleotides, and position of hybridization with crRNA. Among all the oligonucleotides tested, we selected two PS-modified DNA-crRNA (crDuplex11 and crDuplex13), exhibiting the most potent inhibition against Cpf1 expression plasmid (Fig. 1e) for further studies. Recent studies reported that co-delivery of chemically modified Cpf1 mRNA and crRNA improved genome editing efficiency of Cpf1^19^. To further validate the observed inhibitory effects of crDuplex11 and crDuplex13, we delivered crRNA duplexes in the presence of Ψ-modified Cpf1 mRNA in 293T cells. In this case, crDuplex13 exhibited detectable genome editing activity, whereas crDuplex11 was capable of completely disabling Cpf1 function under the same condition (Fig. 1f). Comparing the structures of crDuplex11-crDuplex13, we implied that PS-DNA oligonucleotide with the same length of crRNA (complementary to the whole sequence of crRNA) may be essential to fully inhibit genome editing of Cpf1.

### Switch-off function of PS-DNA oligonucleotides

In the study above, crDuplex11 containing 42-PS linkage modified DNA oligonucleotide (termed ps42DNA) was able to completely switch off the activity of Cpf1 in the hybridization form. Subsequently, we speculate that ps42DNA itself may enable us to inactivate Cpf1 activity and serve as a Cpf1 inhibitor. To test this hypothesis, we separately formulated three components (crRNA targeting *DNMT1* locus, AsCpf1 mRNA, and ps42DNA) using Lipofectamine 3000 reagent, and then simultaneously delivered these three components to 293T cells. Amazingly, the process of genome editing was effectively interrupted when ps42DNA was added together with the other two components (time = 0 h, Fig. 2a). Next, we treated cells with crRNA plus AsCpf1 mRNA followed by the addition of ps42DNA at various time points in order to evaluate inhibition properties of ps42DNA. As shown in Fig. 2a, ps42DNA displayed time-dependent inhibition on genome editing activity. At the 10 h time point, ps42DNA was not able to affect the Cpf1 function. We then investigated dose-dependent inhibitory effects of ps42DNA 5 h post-treatment of CRISPR-Cpf1 components. ps42DNA was found to act in a dose-dependent manner to regulate genomic cleavage (Fig. 2b). These observations indicated that time to add ps42DNA and dose of ps42DNA are two crucial factors to exert switch-off function for this CRISPR-Cpf1 system.

**Fig. 2.**
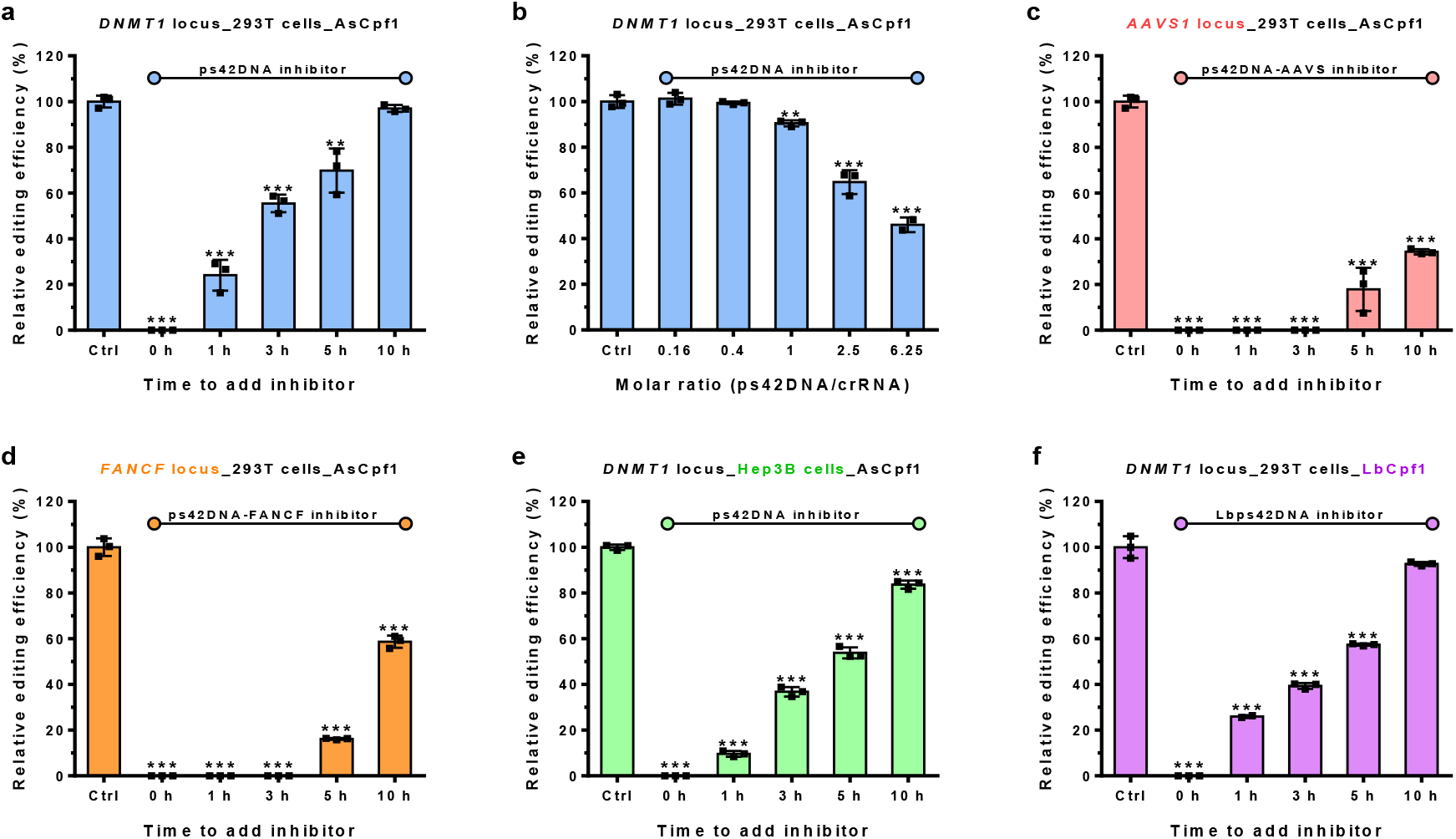
Inhibitory effects of phosphorothioated DNA oligonucleotides on Cpf1-mediated genome editing in human cells. **a** Time-dependent inhibitory effects of ps42DNA on AsCpf1-mediated *DNMT1* genome editing in 293T cells. “Time to add inhibitor”: ps42DNA was added at various time points after treatment with CRISPR-Cpf1 components. **b** Dose-dependent inhibitory effects of ps42DNA on AsCpf1-mediated *DNMT1* genome editing in 293T cells. ps42DNA was added at 5 h after treatment with CRISPR-Cpf1 components. **c** Time-dependent inhibitory effects of ps42DNA-AAVS1 on AsCpf1-mediated genome editing at the *AAVS1* locus in 293T cells. **d** Time-dependent inhibitory effects of ps42DNA-FANCF on AsCpf1-mediated genome editing at the *FANCF* locus in 293T cells. **e** Time-dependent inhibitory effects of ps42DNA on AsCpf1-mediated genome editing at the *DNMT1* locus in Hep3B cells. **f** Time-dependent inhibitory effects of Lbps42DNA on LbCpf1-mediated genome editing at the *DNMT1* locus in 293T cells. Relative genome editing efficiency was determined by the T7E1 cleavage assay, normalized to that of the treatment with wild-type crRNA and AsCpf1 mRNA, and plotted versus time or dose (**, P < 0.01; ***, P < 0.001; two-tailed *t*-test). The corresponding gel images were shown in Supplementary Fig. 2.

### Applicability of ps42DNA in additional gene loci and cell types

To assess the applicability of this approach, we synthesized two additional phosphorothioated oligonucleotides complementary to crRNAs targeting the *AAVS1* and *FANCF* genes (Supplementary Table 1), termed ps42DNA-AAVS 1 and ps42DNA-FANCF, respectively (Supplementary Table 1)^31,32.^ Consistent with the results mentioned above, both ps42DNA-AAVS1 and ps42DNA-FANCF showed time-dependent inhibition of genome editing for their corresponding sequences. Their inhibition potency were higher than that of ps42DNA targeting *DNMT1,* as evidenced by undetectable cleavage at the time points 1, 3 and 5 h (Fig. 2c,d). In addition to 293T cell line, we also evaluated the effects of ps42DNA in Hep3B cells (a human hepatoma cell line). Similarly, ps42DNA showed dramatic inhibition of genome editing in Hep3B cells (Fig. 2e). Next, we examined the same strategy for another Cpf1 ortholog, LbCpf1^1^, using its corresponding phosphorothioated DNA oligo (termed Lbps42DNA, Supplementary Table 1), which exhibited strong inhibition of genome editing in a time-dependent manner similar to ps42DNA (Fig. 2f). Collectively, ps42DNA oligonucleotides are broadly applicable to inhibit Cpf1-mediated genomic cleavage at different gene loci in mammalian cells.

### crDuplexes interact with Cpf1 similar to crRNA

Previous reports showed that Cpf1 and crRNA first formed a binary complex and then exert a conformational change to the Cpf1-crRNA-DNA ternary complex^5^. To study the binary complex of Cpf1 and crDuplexes, we performed an electrophoretic mobility shift assay using AsCpf1 protein. To clarify the complex components, crDuplex11 (hybridized from ps42DNA and crRNA) is termed ps42DNA-crRNA. As a control group, crDuplex2 (hybridized from unmodified DNA and crRNA) is termed uDNA-crRNA in the following assays. As shown in Figure 3, after incubation of crRNA, uDNA-crRNA, or ps42DNA-crRNA with increasing concentrations of AsCpf1 protein, shifted bands occurred comparably in all three cases, reflecting the interactions between crRNA (or crRNA duplexes) and AsCpf1 protein (Fig. 3). Moreover, no single-stranded oligonucleotides in Figure 3 were detected in their corresponding positions. These results indicated that uDNA-crRNA and ps42DNA-crRNA were bound into AsCpf1 in a double-stranded state.

**Fig. 3.**
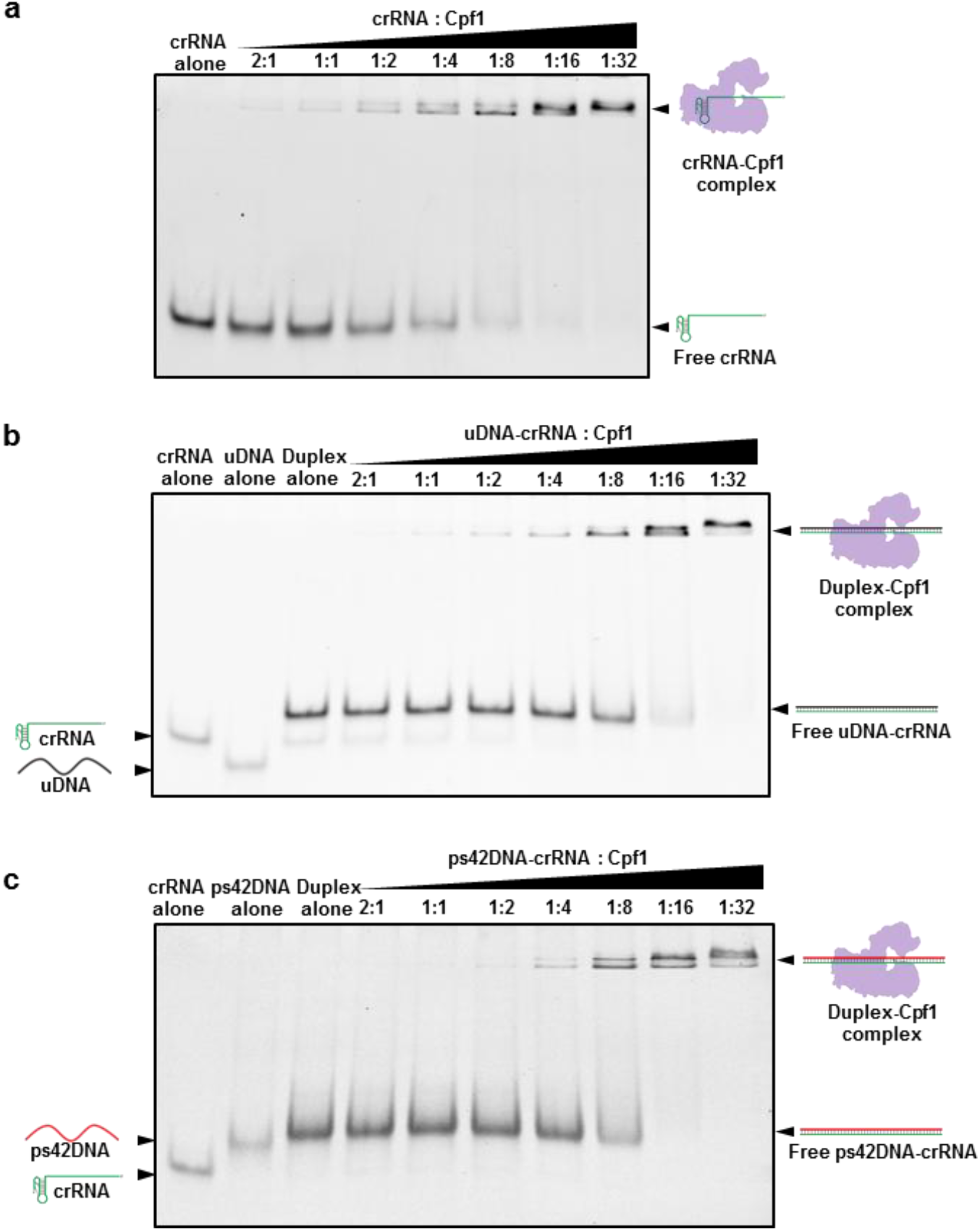
Interactions between Cpf1 and crRNA or crDuplex. Complex formation between AsCpf1 protein and crRNA (**a**), uDNA-crRNA (**b**) or ps42DNA-crRNA (**c**) via electrophoretic mobility shift assays. uDNA, unmodified DNA oligonucleotide.

### Inhibitory effects of ps42DNA in a cell-free condition

Cpf1 utilizes a single-stranded crRNA to recognize and edit target DNA^1,3.^ In order to mimic genome editing in cell studies, the *DNMT1* genomic region was amplified by PCR and used as a double-stranded DNA substrate. We first mixed AsCpf1, crRNA, and DNA substrate to examine the *in vitro* reaction condition. As we increased the amount of AsCpf1, more cleaved fragments were observed (Supplementary Fig. 3a), in accordance with the phenomena found in Figure 3a. To further investigate the inhibitory effects, we added ps42DNA into the reaction mixture at different time points using a fixed molar ratio of 1 : 8 : 1 (crRNA : AsCpf1 : DNA substrate). We found that ps42DNA inactivated AsCpf1 in a time-dependent trend, which was consistent with our observation in the cell studies (Supplementary Fig. 3b). Because this *in vitro* reaction doesn’t need AsCpf1 production, the time frame ranges in minutes other than that in hours range in the cells.

### ps42DNA blocked DNA substrate binding with the complex of Cpf1-crRNA

In order to distinguish different components in the Cpf1 system, we labeled the 3’-end of crRNA with a Cy5 fluorescent probe (termed Cy5crRNA), and then examined its function in 293T cells. Cy5crRNA showed equivalent activity to crRNA, indicating that Cy5 probe is suitable for our mechanism study (Supplementary Fig. 3c). We subsequently carried out the same *in vitro* cleavage assay using Cy5crRNA to visualize the interactions between cleaved DNA fragments and crRNA-Cpf1 complex. As displayed in Figure 4a, the short cleaved fragment (lane 4, white arrow) in the presence of proteinase K was shifted to a higher position (lane 6, white arrow) without treatment of proteinase K under the SYRB Green channel. Meanwhile, we noticed that this shifted band displayed Cy5 fluorescent signal under the Cy5 channel, suggesting the existence of crRNA (Fig. 4a, lane 6 in the middle panel). In Figure 4b, we conducted the reactions with crRNA, uDNA-crRNA, or ps42DNA-crRNA in the absence of proteinase K, and then imaged the gel via both SYRB Green and Coomassie Blue staining (Fig. 4b). The images in Fig 4a and 4b reflect the complex of the crRNA, Cpf1 protein and the short cleaved DNA fragment. As illustrated in Figure 4c, in the presence of proteinase K, three components are released from the complex including crRNA and two cleaved DNA fragments. On the contrary, if no treatment of proteinase K, the short cleaved DNA fragment with intensive interactions with crRNA and Cpf1 is still bounded in the ternary complex. Therefore, this DNA band (short cleaved fragment) was shifted from the position less than 400 bp as shown in the ladder to the position over 500 bp, while the other DNA fragment (long cleaved fragment) remained the same (Fig. 4a,b). Lastly, we quantified the amount of DNA substrate after reactions in Figure 4b by band intensity under the SYRB Green channel in comparison to the same amount of untreated DNA substrate, and found that DNA substrate in both crRNA and uDNA-crRNA groups were much lower than the untreated group, while DNA substrate in the group of ps42DNA-crRNA displayed no significant difference (Fig. 4b,d). These results indicated that ps42DNA hindered the binding of the Cpf1 with the DNA substrate. Taken together, we propose the following mechanism of action (Fig. 4e): Both uDNA-crRNA and ps42DNA-crRNA duplexes were complexed well with Cpf1 protein similar to the wild-type crRNA. After that, the complex of uDNA-crRNA and Cpf1 is able to recognize DNA substrate and induce gene cutting. In the case of ps42DNA, the complex of ps42DNA-crRNA and Cpf1 fully block the binding to DNA substrate, and consequently inactivate the Cpf1 function.

**Fig. 4.**
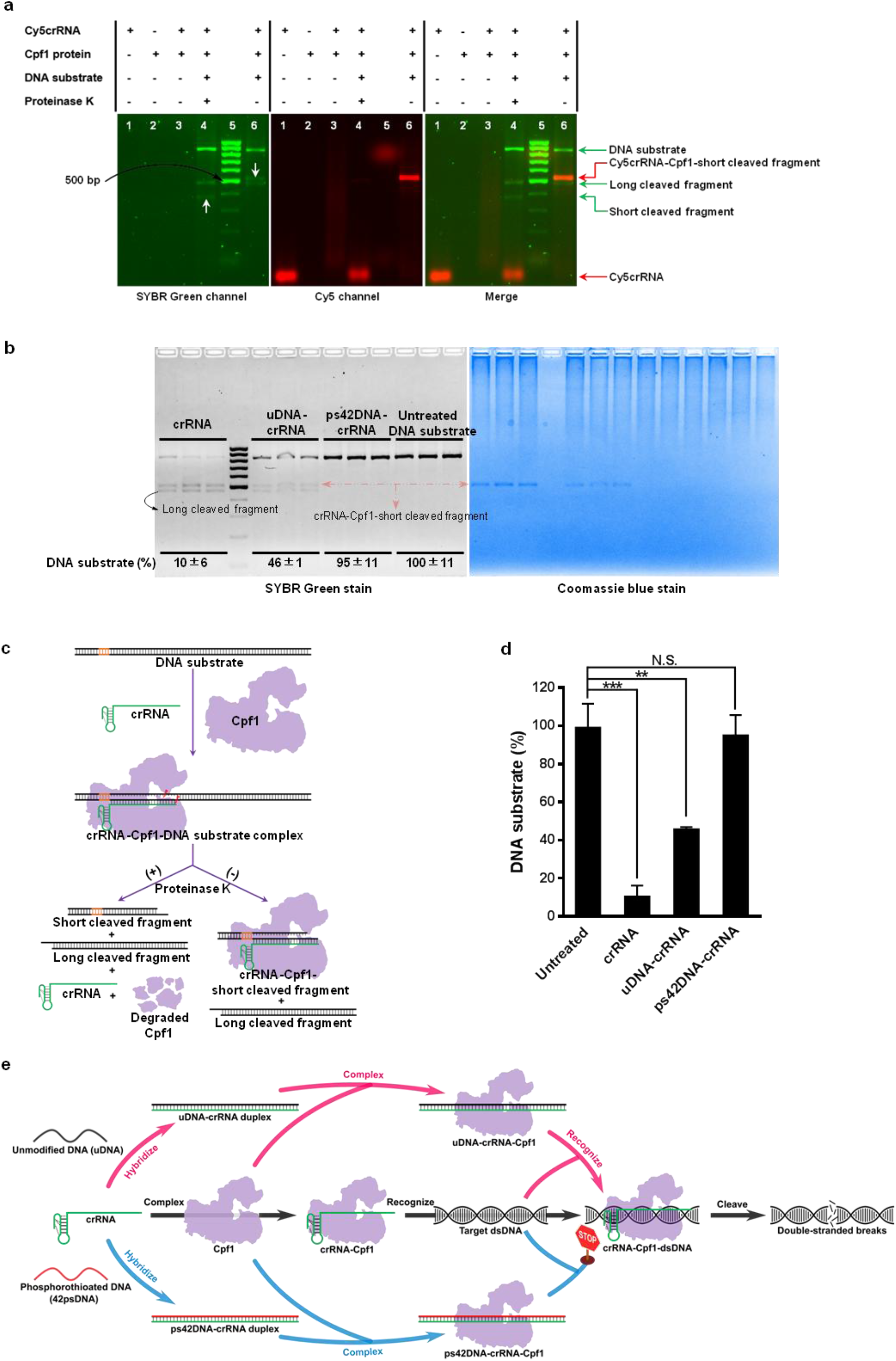
Mechanistic studies for phosphorothioated DNA oligonucleotides on Cpf1-mediated genome editing. **a** Analysis of fluorescently (Cy5) labeled crRNA-mediated *in vitro* cleavage reactions under SYBR (left panel) and Cy5 (middle panel) channel. A merged image is shown in the right panel. **b** Left panel: Quantification of DNA substrate bound with AsCpf1 in the presence of crRNA, uDNA-crRNA or ps42DNA-crRNA, respectively. The uncleaved DNA substrate was quantified by densitometric analysis and normalized to the untreated group. Right panel: Localization of the complex of Cpf1, crRNA and the short cleaved DNA fragment by Coomassie Blue staining. **c** Illustration of Cpf1-mediated *in vitro* cleavage reactions with and without proteinase K digestion. After the formation of crRNA-Cpf1-target DNA ternary complex, DNA substrate is cleaved. With the addition of proteinase K, crRNA and two cleaved DNA fragments are released from the complex. Nevertheless, in the absence of proteinase K, one DNA fragment (long cleaved fragment) without extensive interactions with crRNA and Cpf1 is released form the ternary complex. The remaining ternary complex consists of the other DNA fragment (short cleaved fragment), crRNA, and Cpf1. **d** A histogram of the normalized DNA substrate (%) in **b** (**P < 0.01, ***P<0.001; NS, not significant; two-tailed *t*-test). **e** Proposed mechanism of inhibition for phosphorothioated DNA oligonucleotides. Similar to crRNA, both crRNA-uDNA and crRNA-ps42DNA duplexes were complexed well with the Cpf1 protein. After addition of DNA substrate, the complex of uDNA-crRNA and Cpf1 is able to induce gene cutting, while the complex of ps42DNA-crRNA and Cpf1 block the binding to DNA substrate and inactivate Cpf1 function.

## Discussion

In this study, we designed a series of oligonucleotides in order to regulate the activity of Cpf1. Based on current results, we summarized a list of structure-activity relationships for modulating Cpf1 function: (I) In general, unmodified DNA and RNA oligonucleotides showed moderate effects on Cpf1-mediated genome editing activity regardless of the hybrid region and the length of oligonucleotides tested. (II) 2’-fluoro-modified oligonucleotides substantially interfered the Cpf1-mediated cleavage. (III) 2’-0-methyl-modified oligonucleotides maintained the performance of the wild-type crRNA. (IV) PS-modified DNA, partially complementary to crRNA inhibited Cpf1 activity to some extent. (V) PS-modified DNA rather than PS-modified RNA, fully complementary to crRNA was able to inactivate Cpf1 function in a time-and dose-dependent manner. Most importantly, we discovered that phosphorothioate (ps)-modified DNA oligonucleotides are effective inhibitors for the Cpf1. These PS-modified DNA oligonucleotides enabled us to switch off genome editing activities of both AsCpf1 and LbCpf1. Also, similar phenomenon was observed in three genomic loci. The inhibitory effect is time-and dose-dependent in human cells.

Electrophoretic mobility shift assays showed that AsCpf1 protein bound to both single-stranded crRNA and double-stranded crRNA duplexes. These results indicated that oligonucleotides may not affect Cpf1 protein binding. Both Cy5crRNA and uDNA-Cy5crRNA formed a ternary complex with one of the cleaved DNA fragments and Cpf1, while ps42DNA-Cy5crRNA didn’t. Previous studies reported that a conformational change occurs from the Cpf1-crRNA binary state to the Cpf1-crRNA-DNA ternary complex^5^. This structural transition facilitates the seed segment of crRNA developing an A-form conformation in order to pair with target DNA and thereby induce gene cutting^5^. We speculate that PS-modified DNA oligonucleotides may hinder the binding between crRNA-Cpf1 complex and target DNA. Overall, these results provide new tools to further understand and modulate the CRISPR-Cpf1 system. In case that acute toxic effects of the CRISPR system occur in clinical use, this strategy may sever as an important antidote.

## Methods

### Preparation of crRNA duplexes

crRNAs (Supplementary Fig. 1) were synthesized by TriLink BioTechnologies via a solid-phase DNA/RNA synthesizer, purified by polyacrylamide gel electrophoresis system, and characterized by electrospray-ionization mass spectrometry. Unmodified oligonucleotides were obtained from Eurofins Genomics, and chemically modified oligonucleotides were obtained from Integrated DNA Technologies. crRNA duplexes (crDuplex1-21, Supplementary Fig. 1) were generated by hybridization of an equivalent molar of AsCpf1 crRNA targeting *DNMT1* and the customized oligonucleotides (Supplementary Table 1). The mixture was heated to 95 °C for 30 s in Tris-EDTA buffer, followed by gradient cooling to room temperate at a rate of 0.1 °C/s. crRNA duplexes derived from AsCpf1 crRNAs targeting *AAVS,* and *FANCF* loci, and LbCpf1 crRNAs targeting *DNMT1* locus were obtained through the same procedure. The sequences of all oligonucleotides used were listed in Supplementary Table 1.

### Genome editing induced by crRNA duplexes

Human 293T and Hep3B cells are from American Type Culture Collection. Cells were seeded in 24-well plates in medium (Dulbecco’s Modified Eagle’s Medium for 293T cells, and Eagle’s Minimum Essential Medium for Hep3B cells) supplement with 10% FBS at a density of 70,000∼100,000 cells per well for 24 h. Cells were then treated with 38 or 114 pmol crRNA variants formulated with Lipofectamine 3000 (Life Technologies) in Opti-MEM I reduced serum medium following the manufacturer’s instructions. Meanwhile, 500 or 1500 ng of Cpf1 plasmid or mRNA (TriLink BioTechnologies) were formulated with the same protocol and added to each well. Cells treated with the wild-type crRNA plus Cpf1 plasmid (or Cpf1 mRNA) served as the control group.

### Time-and dose-dependent inhibition of Cpf1 activity

In order to study the effects of time interval, 2.5 times molar excess of PS-modified DNA over wild-type crRNA was added to cells treated with the wild-type crRNA plus Cpf1 mRNA at different time points (0, 1, 3, 5 and 10 h). In the case of dose-dependence assay, crRNA and Cpf1 mRNA-treated cells were exposed to different molar of PS-modified DNA (the molar ratio of PS-DNA : crRNA ranged from 1 : 6.25 to 6.25 : 1) at 5 h. In both conditions, the end point was 48 h after the addition of a combination of wild-type crRNA and Cpf1 mRNA.

### T7E1 enzymatic cleavage assays

Two days after treatment, genomic DNA was harvested from treated cells using the DNeasy Blood & Tissue Kit (QIAGEN). Polymerase chain reactions (PCRs) were then performed using Q5 Hot-start High-Fidelity DNA Polymerase (New England Biolabs). Primers (Eurofins Genomics) flanking the targeted region were listed in Supplementary Table 2. The PCR products were then annealed in NEBuffer 2 (50 mM NaCl, 10 mM Tris-HCl, 10 mM MgCl_2_, 1 mM DTT, New England Biolabs) and subsequently digested by T7 Endonuclease I (T7E1, New England Biolabs) at 37 °C for 30 min. The fraction cleaved was separated on 2% agarose gels, visualized on the ChemiDoc MP imaging system (Bio-Rad Laboratories), and analyzed by the Image Lab 5.2 analysis software (Bio-Rad Laboratories).

### Electrophoretic mobility shift assays

Electrophoretic mobility shift assays were performed using 0.5 pmol of wild-type crRNA (or its variants) and 0.25 pmol∼16 pmol of AsCpf1 protein (generously provided by New England Biolabs) at 37 °C for 30 min in 7 uL of NEBuffer 3 (100 mM NaCl, 50 mM Tris-HCl, 10 mM MgCl_2_, 1 mM DTT, New England Biolabs). The binding reactions were fractionated on 15% non-denaturing TBE polyacrylamide gels (Bio-Rad Laboratories), stained by SYBR Gold dye (Thermo Fisher Scientific), and detected on the ChemiDoc MP imaging system.

### Gene editing in a cell-free condition

To mimic genomic cleavage in cells, the target *DNMT1* locus was amplified from genomic DNA isolated from untreated 293T cells using Q5 High-Fidelity DNA Polymerase and primers listed in Supplementary Table 2. PCR amplicons used as the DNA substrate were cleaned up with QIAquick PCR Purification Kit (QIAGEN). Prior to cleavage, crRNA (0.5 pmol) was equilibrated with different concentrations of AsCpf1 protein (crRNA: AsCpf1 is from 2 : 1 to 1 : 16) for 30 min at 37 °C in 7 uL of NEBuffer 3 solution. Subsequently, 3 uL of PCR amplicons (0.5 pmol, in NEBuffer 3 solution) was incubated with the above complex at 37 °C for 30 min. The reactions were terminated by adding 1 uL of proteinase K, resolved on 2% agarose gels, visualized by EZ-Vision In-Gel staining, and imaged on the ChemiDoc MP imaging system. To character *in vitro* inhibition of Cpf1-mediated cleavage activity by ps42DNA, crRNA, AsCpf1 protein, and DNA substrate were incubated at 37 °C for 30 min at the molar ratio of 1 : 8 : 1 (crRNA : AsCpf1 protein: DNA substrate). At different time points (t = 0, 5, 10, 15, 20, 25 min), 2.5-fold molar excess of ps42DNA over crRNA was added to the reaction mixture. The remaining procedures were the same as described above.

### Visualization of crRNA, Cpf1 protein, and DNA substrate using multiple imaging channels

To assess the effects of fluorescence labeled crRNA on genome editing efficiency, Cy5-labeled crRNA (Cy5crRNA, Integrated DNA Technologies) was first evaluated in 293T cells as described above. To visualize the guide RNA, crRNA *in vitro,* Cy5crRNA was complexed with Cpf1 protein at 37 °C for 30 min at a molar ratio of 1 : 8, followed by incubation with an equivalent mole of DNA substrate at 37 °C for 30 min. The reaction mixture was digested with or without proteinase K prior to agarose gel electrophoresis, and the signals of reaction products were collected via a ChemiDoc MP imaging system under the SYBR Green and Cy5 channels. To image the Cpf1 protein, crRNA, uDNA-crRNA or ps42DNA-crRNA was complexed with Cpf1 protein at 37 °C for 30 min at a molar ratio of 1 : 8. The remaining procedures were the same as mentioned above, except that the agarose gel was stained with Coomassie Blue and imaged under the SYBR Green and Coomassie Blue channels. The uncleaved DNA substrate was quantified by densitometric analysis and normalized to that of the untreated DNA substrate.

## Author Contributions

B.L. designed and performed all experiments, analyzed data and wrote the manuscript. C.Z. designed and performed experiments related to in vitro cleavage assay. X.Z. designed and conducted fluorescence analysis. W.L., X.L., W.Z., and C.Z. conducted cell analysis. Y.D. conceived and supervised the project and wrote the manuscript. The final manuscript was edited and approved by all authors.

**Supplementary information** is available for this paper.

